# A sub-exponential branching process to study early epidemic dynamics with application to Ebola

**DOI:** 10.1101/797878

**Authors:** Alexander E. Zarebski, Robert Moss, James M. McCaw

## Abstract

Exponential growth is a mathematically convenient model for the early stages of an outbreak of an infectious disease. However, for many pathogens (such as Ebola virus) the initial rate of transmission may be sub-exponential, even before transmission is affected by depletion of susceptible individuals.

We present a stochastic multi-scale model capable of representing sub-exponential transmission: an in-homogeneous branching process extending the generalised growth model. To validate the model, we fit it to data from the Ebola epidemic in West Africa (2014–2016). We demonstrate how a branching process can be fit to both time series of confirmed cases and chains of infection derived from contact tracing. Our estimates of the parameters suggest transmission of Ebola virus was sub-exponential during this epidemic. Both the time series data and the chains of infections lead to consistent parameter estimates. Differences in the data sets meant consistent estimates were not a foregone conclusion. Finally, we use a simulation study to investigate the properties of our methodology. In particular, we examine the extent to which the estimates obtained from time series data and those obtained from chains of infection data agree.

Our method, based on a simple branching process, is well suited to real-time analysis of data collected during contact tracing. Identifying the characteristic early growth dynamics (exponential or sub-exponential), including an estimate of uncertainty, during the first phase of an epidemic should prove a useful tool for preliminary outbreak investigations.

**Author Summary:** Epidemic forecasts have the potential to support public health decision making in outbreak scenarios for diseases such as Ebola and influenza. In particular, reliable predictions of future incidence data may guide surveillance and intervention responses. Existing methods for producing forecasts, based upon mechanistic transmission models, often make an implicit assumption that growth is exponential, at least while susceptible depletion remains negligible. However, empirical studies suggest that many infectious disease outbreaks display sub-exponential growth early in the epidemic. Here we introduce a mechanistic model of early epidemic growth that allows for sub-exponential growth in incidence. We demonstrate how the model can be applied to the types of data that are typically available in (near) real-time, including time series data on incidence as well as individual-level case series and chains of transmission data. We apply our methods to publically available data from the 2014–2016 West Africa Ebola epidemic and demonstrate that early epidemic growth was sub-exponential. We also investigate the statistical properties of our model through a simulation re-estimation study to identify it performance characteristics and avenues for further methodological research.

## 1. Introduction

Physical systems can rarely support exponential growth for extended periods; during an epidemic, depletion of susceptible individuals leads to reduced transmission and, if intervention measures have not already done so, cause incidence to decline. Despite recent work showing the initial transmission of many diseases is sub-exponential, it is still common to see epidemics represented by models in which transmission grows exponentially Viboud et al. (2016). This is concerning because exponential growth is extremely sensitive to its growth rate parameter, which can inflate the variance of forecasts. During an outbreak of a novel pathogen, uncertainty in the growth rate is almost guaranteed. Furthermore, the likely impact of an intervention, such as social-distancing or deployment of a vaccine, is likely to be highly sensitive to the estimate for the growth rate parameter.

The quantitative models used in epidemiology vary, from simple phenomenological models Lega and Brown (2016); Nouvellet et al. (2018) to complex agent-based simulations Ajelli et al. (2016); Merler et al. (2015).

Typically, the simpler phenomenological models — while able to produce exponential or sub-exponential growth — lack the mechanistic underpinning to answer relevant question (e.g., what will be the effect of vaccinating 20% of the population?) and so have arguably limited application in outbreak investigations. At the other end of the complexity spectrum, agent-based models, with their high biological fidelity, allow for conceptually simple explorations of the impact of interventions. However, there is a cost: their complexity makes them difficult to reason about mathematically. They are also computationally intensive, making statistical analysis and so assessment of the early growth characteristics and potential impact of interventions, challenging.

Here, in the context of early outbreak investigation, we demonstrate how an inhomogeneous branching process formulation can overcome some of the challenges described above: the mismatch between exponential growth of transmission and observations, and the difficulty of finding a model with a mechanistic basis which is still mathematically tractable. A temporal in-homogeneity in the branching process ensures the generation sizes grow algebraically (in expectation), instead of the typical geometric/exponential growth. Branching processes can be viewed as either a tree, where it describes who-infected-whom, or as a time series to describe the total number of cases through time. As such, they are a good example of a multi-scale model. Unlike many of the complex mechanistic models, the simplicity of the branching process means it is possible to reason about them quantitatively and work with them computationally.

We explore the use and properties of this model from three perspectives. First, we use the branching process in a hierarchical model of transmission of Ebola virus in West Africa. Using publicly available data made from the World Health Organisation we demonstrate how the branching process can faithfully describe observed epidemics. Second, we fit the branching process to two different types of data: chains of infection and time series of cases of Ebola virus disease (EVD) from the West African Ebola epidemic (2014–2016). Our analysis demonstrates the model provides broadly consistent parameter estimates using either data type, despite differences between the data sets. While the sub-exponential transmission of Ebola virus has been previously noted, Chowell et al. (2015), the branching process allows us to go further, supporting this claim through the interrogation of a new data set: a fully resolved infection tree inferred by Faye *et al* Faye et al. (2015). Third, to investigate the extent to which one might expect the previous result (i.e., obtaining similar estimates from each data type) to generalise, we performed a simulation study. The goal of this simulation study was not to investigate the utility of each data type for estimating the parameters *per se*, but to ask whether or not both data types, when derived from the same epidemic, produce concordant estimates.

## 2. Methods —— Model and analysis

We derive the branching process in terms of a generic cumulative incidence function, i.e., a function describing the total number of cases that have occurred by a given time. We then consider the special case of a cumulative incidence function previously used to analyse time series of Ebola in West Africa Viboud et al. (2016). Finally, we construct a likelihood function for this model, both in terms of a time series of cases and for observations of the number of secondary cases generated by individuals.

### 2.1. Construction of the in-homogeneous branching process model

Let 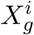 denote the number of secondary infections due to individual *i* in generation *g* and *Z_g_* the total number of infectious individuals in that generation, i.e., the sum of the 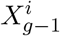. We derive an in-homogeneous branching process where the *expected* generation sizes are 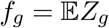.

Usually, the expected number of infectious individuals in a branching process grows exponentially/geometrically in the number of generations of transmission. For example, if 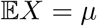 then 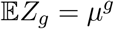. The branching process derived below has expected generation sizes (i.e., the 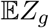) which can follow any given monotonically increasing function. The notation used in this construction is summarised in Table 1.

**Table 1.**
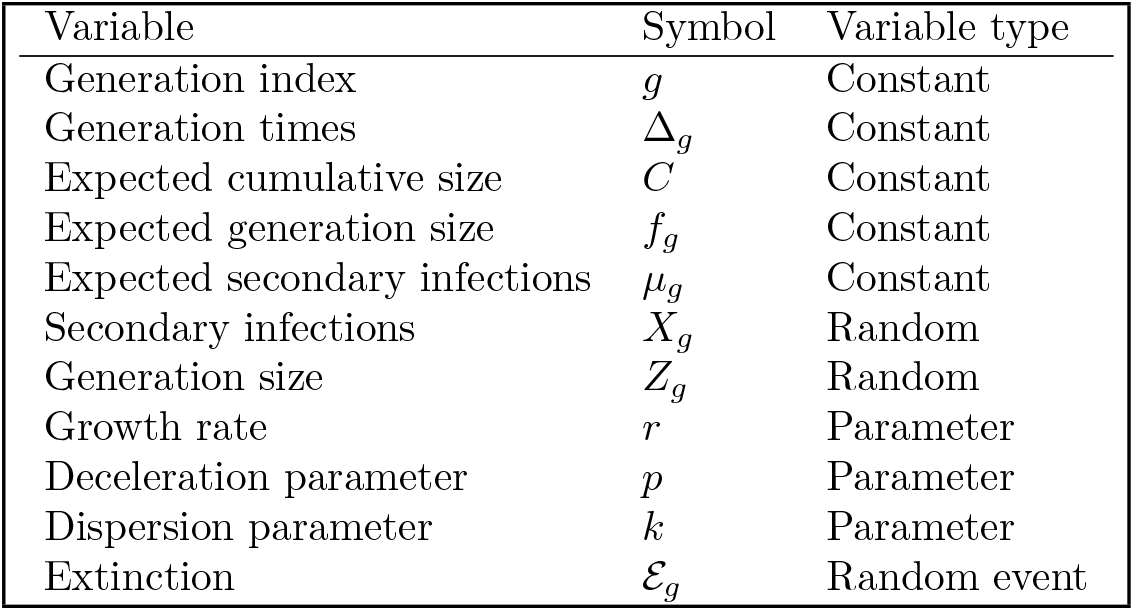
Notation used for the branching process.

Let *C*(*t*) be the *expected* cumulative incidence by time *t*, i.e., the number of infections we would expect to occur by time *t*. Evaluated at multiples of the serial interval, *C* yields the generation sizes, *f_g_* for *g* = 1, 2, …

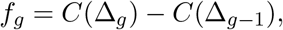

where Δ_*g*_ is the time of the *g*th generation. The first value of this sequence is *f*_0_ = *Z*_0_, the number of infectious individuals in the first generation. Then, assuming the 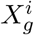 are independent with mean *μ_g_* = *f*_*g*+1_/*f_g_* we observe

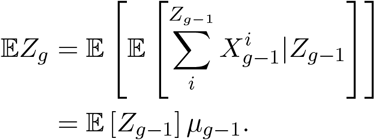

The solution to this recurrence is

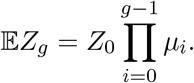

So 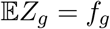 from the definition of *μ_g_*.

In summary, by fixing the expected value of the offspring distribution (in terms of the generation) we obtain a branching process which, on average, has an expected cumulative incidence *C*. This construction enables us to capture the behaviour of a phenomenological model which is known to fit observations better than exponential/geometric growth, while maintaining a mechanistic foundation because it explicitly represents the individuals in the population.

### 2.2. The cumulative incidence function

The construction above assumes a cumulative incidence function, *C*. We use the generalized growth model Viboud et al. (2016) defined by

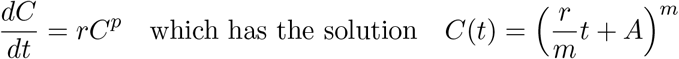

where *m* = 1/(1 − *p*) and 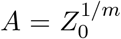, with initial condition *C*(0) = *Z*_0_. The growth rate, *r*, is as for standard exponential growth. The generalisation enters through the inclusion of the exponent *p*.

The parameter *p* is referred to as the *deceleration parameter*; it influences the dynamics of transmission. For 0 < *p* < 1 the incidence interpolates through polynomials limiting to exponential growth as *p* → 1. For *p* < 1 there is a diminishing increase in the force of infection with each additional infection. When *p* = 0 the force of infection is constant, for *p* = 1/2 (when m = 2) the incidence grows linearly (since the incidence is the derivative of the cumulative incidence by definition), *p* = 2/3 provides quadratic growth and with *p* = 1 we recover exponential growth in incidence.

Previous analyses suggest that the spread of diseases, such as Measles, HIV/AIDS, and FMD, are well explained by values of *p* < 1; 0.51 (0.47, 0.55), 0.5 (0.47, 0.54), and 0.42 (0.27, 0.58) respectively Viboud et al. (2016).

### 2.3. The offspring distribution

Since epidemiological count data is frequently over-dispersed (with respect to the Poisson distribution) we use the negative binomial distribution for the offspring distribution. Over-dispersion in count data can occur for many reasons Lindén and Mäntyniemi(2011), for case counts in an epidemic, *superspreaders* can play an important role Lloyd-Smith et al. (2005). We parameterise the negative binomial in terms of its mean, *μ*, and a shape parameter, *k*, (a.k.a. the “dispersion parameter”). Under this parameterisation the variance, *σ*^2^, grows quadratically in *μ*

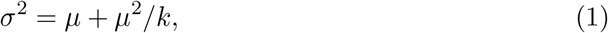

so as *k* → ∞ we recover the Poisson distribution. Since the mean value is determined by the cumulative incidence function (*μ* = *C*) this choice of offspring distribution only introduces a single additional parameter, *k*.

### 2.4. Likelihood function for time series and chains of infection

Early work by Wald in the 1940’s demonstrated the importance of survivorship bias. The importance of subtleties in the provenance of data, and how to account for this via conditioning is well understood in phylogenetics Stadler (2013) yet does not appear to have permeated to the same degree into the epidemiology literature (notable exceptions being the work of Mercer Mercer et al. (2011) and Rida Rida (1991)).

Popular estimators of the basic reproduction number, *R*_0_, are biased towards *overestimation* in the early stages of an epidemic, Mercer et al. (2011). We condition the process against extinction in the likelihood during fitting to mitigate this bias. In short — by virtue of being observed — the outbreak must have avoided stochastic extinction Rida (1991).

Realisations from the branching process are naturally viewed as a tree, with the edges indicating who infected whom. However, as the notation suggests, this process can also be viewed as a sequence of generation sizes, *Z*_0:*g*_. We refer to this representation of the process as the *population view*. As we will see, the ability to represent a process as both a tree and a time series is very useful when making use of multiple data types.

First we consider the likelihood function for the time series data, conditioned against extinction over the observed generations. We extend the notation introduced in section 2.1 to specify the (geographic) location (denoted by *j*): 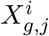 denotes the number of infections caused by the *i*th member of the *g*th generation in location *j*, and *Z_g,j_* denotes the number of cases in generation *g* in location *j*. For ease of notation, we will often drop the indices where they are clear by context.

By definition *X_g_* has a negative binomial distribution with mean μg and shape parameter *k*, so the moment-generating function (MGF) is

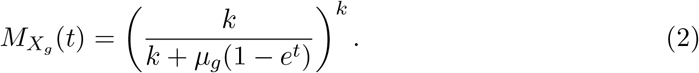

Since *Z_g_*|*Z*_*g*−1_ is the sum of *Z*_*g*−1_ independent *X*_*g*−1_, the MGF is given by the product

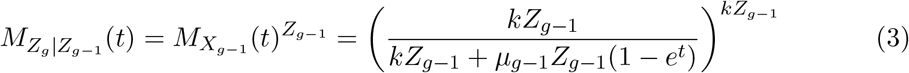

and hence *Z_g_*|*Z*_*g*−1_ is also negative binomial with mean *μ*_*g*−1_*Z*_*g*−1_ and shape parameter *kZ*_*g*−1_. Since the generation sizes form a Markov chain and each location is assumed to have an independent epidemic, the likelihood of all of the time series data is

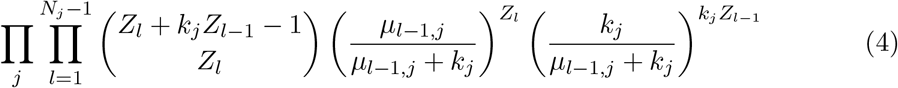

where *N_j_* denotes the number of generations that were observed in location *j*.

In order to condition the process against extinction during the observed generations we use the probability that the process is extinct by the time of the last observation at that location. To compute this probability, we consider the probability-generating function (PGF) of the generation sizes, *G_Z_g__*(*t*). Since we are working with the PGF (rather than the MGF) the sum of *Z*_*g*−1_ independent *X*_*g*−1_ leads to the composition

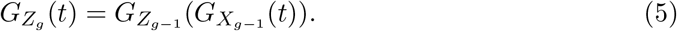

Iterating this *g* times, and noting that *G*_*Z*_0__(*t*) = *t*^*Z*_0_^ leads to

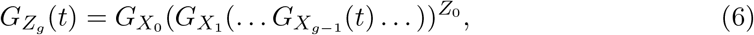

where *G_X_g__*(*t*) = (*k*/(*k* + *μ*(1 − *t*)))^*k*^. The composition produces a complicated expression, but for moderate *g* this is not an issue computationally. The probability of extinction is the zero-th order coefficient of the PGF, hence the probability of extinction by generation *g* is *G_Z_g__*(0).

Putting the previous results together, we obtain the conditional likelihood for the observed times series:

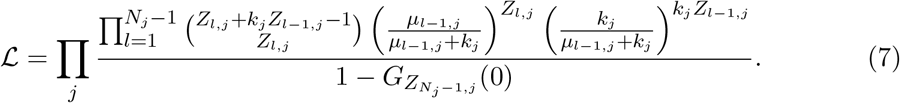

For secondary infections data, the probability of extinction conditional upon partial observations is prohibitively expensive to evaluate since it requires integrating over all the possible hidden infection trees. Subsequently, when working with secondary infections data we do not condition the process against extinction. Instead we treat each count of secondary infections as an independent sample from the offspring distribution. The likelihood is

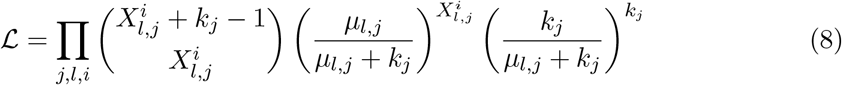

We conclude this section with a few remarks on computation. When computing with the expressions above for the likelihoods, the log-likelihood is used to avoid under-flow/overflow issues. Moreover, the use of a probabilistic programming language (such as Stan as used here) will handle this expression and its gradient in a numerically stable way. Therefore in practice, beyond specifying the process as a graphical model, the only requirement is to implement the computation of the extinction probability.

### 2.5. Epidemiological data from the West African Ebola epidemic 2014–2016

Data of cases of EVD in Guinea, Liberia and Sierra Leone from 2014–2016 were obtained from the WHO World Health Organization (2018). We extracted confirmed cases from the patient data and then selected the longest stretch of consecutive weeks (the temporal resolution of the data) in which there was at least one confirmed case for each country. This process was repeated to generate a time series for each of the countries considered. The longest stretches occurred at the beginning of the epidemic for both Guinea and Sierra Leone, while several isolated cases were removed from the start of the Liberian time series. These time series were aggregated by fortnight as a proxy for generations of transmission since the Ebola virus has an approximate 14 day generation time Chowell and Nishiura (2014). The first 20% of cases were used in the analysis (as the *Z*_0:*G*_) to represent transmission during the initial stage of the epidemic.

The WHO data also includes approximate locations for each case. Using this information, we extracted another time series specific to Conakry, the capital city of Guinea.

Faye *et al* Faye et al. (2015) resolved an infection tree for cases from Conakry and the towns of Boffa and Telimele resulting in the data shown in Figure 1. Of the 193 confirmed and probable cases reported from these locations, 152 were placed in the tree with 106 of these from Conakry. To avoid the effects of re-importation we only used cases from Conakry that were not re-introductions from Boffa or Telimele, leaving 98 cases in the tree.

**Fig. 1.**
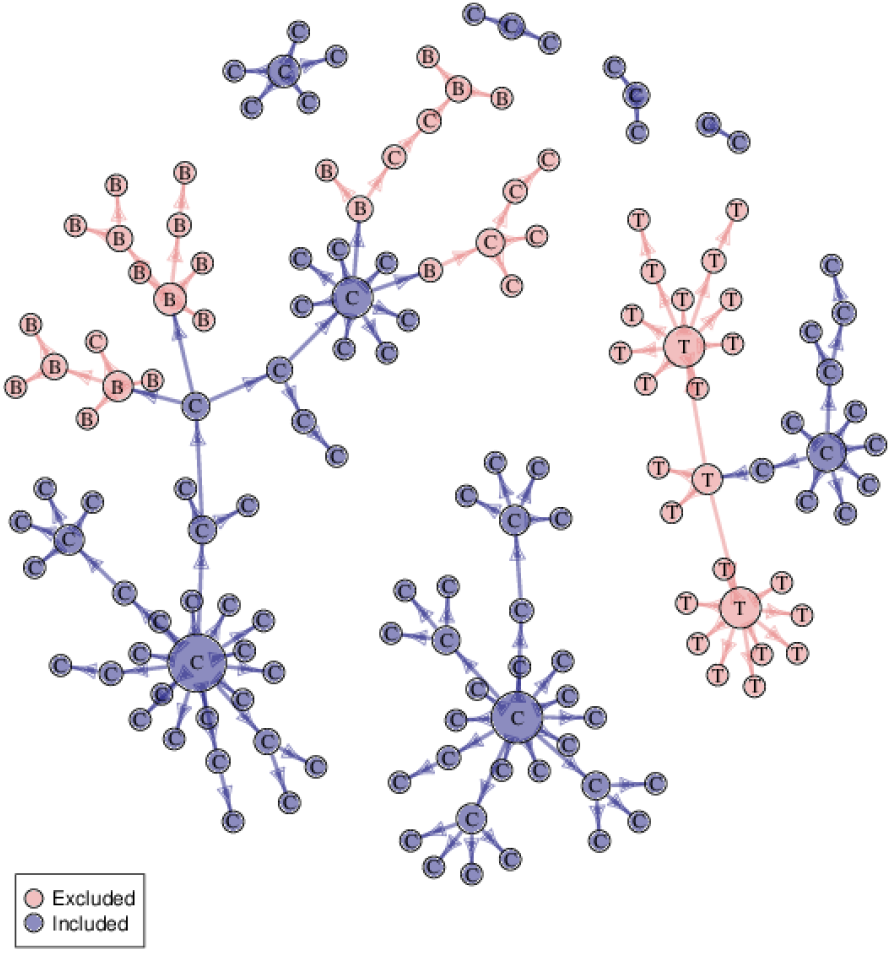
The infection tree from Faye et al Faye et al. (2015). The colour of the nodes indicates whether the data were included in the analysis and the labels indicate where the infection occurred.

In the case of Conakry, it is important to note that there are important differences between the data sets. The time series is specific to confirmed cases while the tree contains both confirmed and probable cases. And the number of cases in the time series is far greater than the number in the infection tree.

### 2.6. Inference method for the time series model

The confirmed cases of EVD in the three West African countries were modelled as time series of generation sizes using the population-level formulation of the branching process. We considered a hierarchical model in which the model parameters for each country are drawn from a common prior distribution which is also estimated. The prior distributions used for the parameters in this model are shown in Table 2. We computed the marginal prior distributions of the model parameters numerically to visually inspect the difference between the prior and posterior distributions.

**Table 2.** Prior distributions used for the model parameters in the hierarchical model described in Section 2.6.

| Type | Parameter | Prior |
| --- | --- | --- |
| Hyperparameter | $\alpha_p$ | Uniform(1, 5) |
| | $\beta_p$ | Uniform(1, 5) |
| | $\mu_r$ | Normal(0, 1) |
| | $\mu_k$ | Normal(0, 1) |
| Parameter | $p$ | Beta($\alpha_p, \beta_p$) |
| | $r$ | Lognormal($\mu_r, \sigma^2$) |
| | $k$ | Lognormal($\mu_k, \sigma^2$) |
| Constant | $\sigma^2$ | 1/6 |

The model was implemented in Stan Carpenter et al. (2017) and Hamiltonian Monte Carlo (HMC) was used to sample from the posterior distribution. Four HMC chains were run; the first 1000 samples of each chain were discarded as burn-in before a further 5000 samples were taken. Of the 5000, this was thinned by a factor of 5 to obtain the final 1000 samples for each chain. The chains appeared to have converged and mixed well: this was established via visual inspection and the 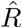-statistic (< 1.01 for all variables). The effective sample size was appropriate given the dimensionality of the problem: for all variables in excess of 80% of the full number of iterations. Subsequently, the posterior samples where taken to provide a good representation of the posterior distribution.

### 2.7. Comparison of time series and chain of infection data from Conakry (Guinea)

As shown in Section 2.4, the branching process can be viewed at the individual or population scale. This prompts the question of whether data collected at each of these scales is equally informative about the parameters of the process, i.e., whether there is any advantage one over the other. We consider two data sets collected in Conakry (the capital city of Guinea) from the Ebola epidemic of 2014–2016: a time series of the number of confirmed cases each week (population scale data), and an infection tree describing who infected whom in a subset of cases (individual scale data). We fit the branching process to both data sets in order to determine whether they would lead to concordant parameter estimates. Note, similarity of the estimates was not guaranteed *a priori*, since while they are both observations of the same epidemic, the data sets consist of different cases. The time series has all the confirmed cases from Conakry, while the infection tree contains only a subset of the confirmed cases but it also contains suspected cases which were excluded from the time series Faye et al. (2015).

We used the population view of the branching process to model the time series of confirmed cases from Conakry. For the secondary infections tree from Conakry (described in Section 2.5), we modelled the number of secondary infections from each individual as an independent sample from the offspring distribution. This takes the form of pairs, 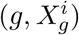, one for each individual, where *g* is their infection generation (the node’s depth in the tree) and 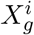 is their number of secondary infections (the out-degree of the node).

The prior distribution used is shown in Table 3. Fitting the model to each data set allows us to investigate whether these views of the same epidemic are consistent.

**Table 3.** Prior distributions used for the model parameters in the comparison of the two data types from Conakry.

| Type | Parameter | Prior |
| --- | --- | --- |
| Parameter | $p$ | Beta(1.5, 1.5) |
| | $r$ | Lognormal(1, $\sigma^2$) |
| | $k$ | Lognormal(1, $\sigma^2$) |
| Constant | $\sigma^2$ | 1/6 |

The models were implemented in Stan and Hamiltonian Monte Carlo (HMC) was used to sample from the posterior distribution. Four HMC chains were run; the first 10000 samples of each chain were discarded as burn-in before a further 10000 samples were taken. Of the 10000, this was thinned by a factor of 10 to obtain the final 1000 samples for each chain. The chains appeared to have converged and mixed well: this was established via visual inspection and the R-statistic (< 1.01 for all variables, with most < 1.001). The effective sample size was sufficiently large, in excess of 90% of the true sample size for all variables. Subsequently, the posterior samples where taken to provide a good representation of the posterior distribution.

### 2.8. Simulation re-estimation study

We carried out a simulation study to investigate whether estimates derived from time series and secondary infections data are concordant and how this depends on the number of secondary infections observed. The goal of this study was to determine the regularity with which the estimates agree, rather than the accuracy with which they capture the dynamics of the epidemic. We simulated a Reed-Frost (RF) epidemic model 1000 times, recording who-infected-whom in each generation Brauer et al. (2008), as described in Supporting Materials. Note that the RF-model assumes a finite population while the branching process implicitly assumes an infinite population. Subsequently, in the RF-model the susceptible pool can be depleted during the epidemic – retarding transmission – eventually causing incidence to decline to zero. In addition to allowing us to investigate agreement between the estimates, fitting the branching process to realisations of the RF-model demonstrates how the model handles deviations from the assumptions used in its construction.

The models were implemented in Stan and L-BFGS was used to approximate the maximum a posteriori probability (MAP) for each of the simulations. Due to the large number of replications considered it was not feasible to check the output of each optimisation manually, instead it was left to the implementation of the optimisation algorithm to determine whether the computation had converged or whether a numerical issue had been encountered (in which case the simulation and optimisation were repeated).

## 3. Results

### 3.1. Hierarchical model fit of the in-homogeneous branching-process model to the EVD data

Figure 2 shows the fit of the hierarchical model to time series of confirmed cases of EVD from Guinea, Liberia and Sierra Leone. The credible intervals on the figures show the uncertainty in the *expected incidence*, i.e., the 50% and 95% credible intervals for 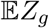.

**Fig. 2.**
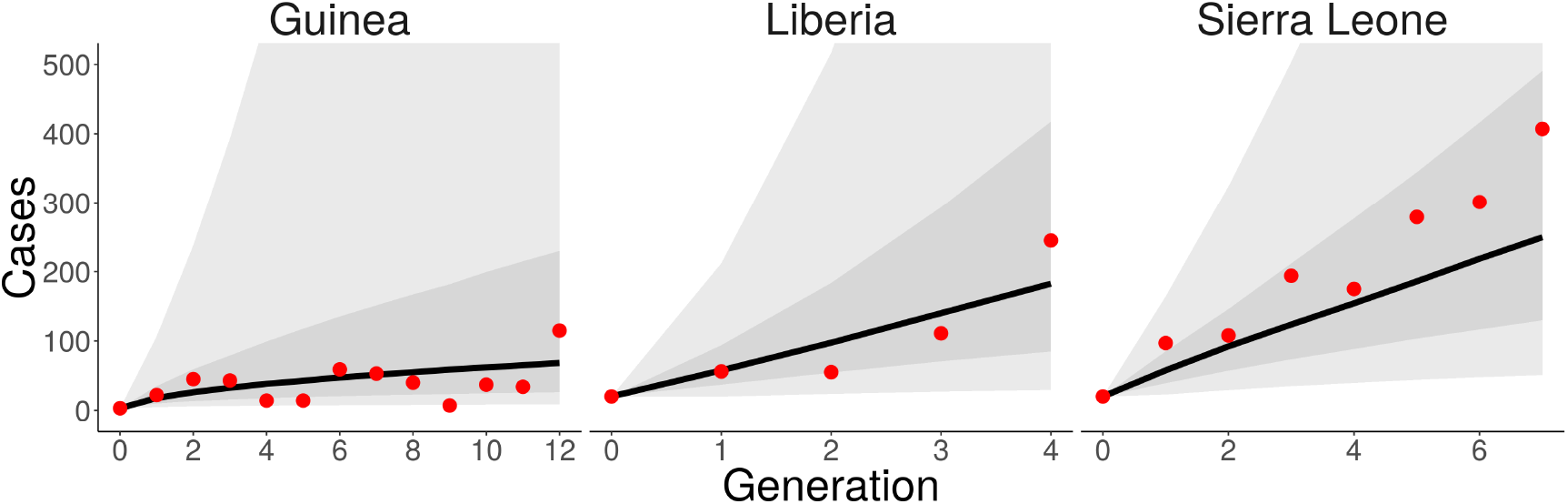
The branching process fit to time series of confirmed cases of EVD from Guinea, Liberia and Sierra Leone. The expected generation sizes (the model fit) are shown as a solid line with the 50% and 95% credible interval on this estimate shown as a grey ribbon. The observed case counts are shown as red points.

Figure 3 shows the marginal posterior distributions of the *logarithm* of the growth rate, the deceleration parameter and the *logarithm* of the dispersion parameter respectively. Figure 3b shows the posterior mass for *p* has accumulated around 0.5 for all three countries; in the model, this corresponds to approximately linear growth in the incidence. Another way to view this would be that the cumulative incidence had quadratic growth. Recall from Equation (1) that the variance scales with the inverse of the dispersion parameter. For each of the countries the dispersion parameter, *k*, has converged to small values indicating that the variance scales quickly with the mean incidence. This suggests stochasticity played an important role in the initial transmission in these countries.

**Fig. 3.**
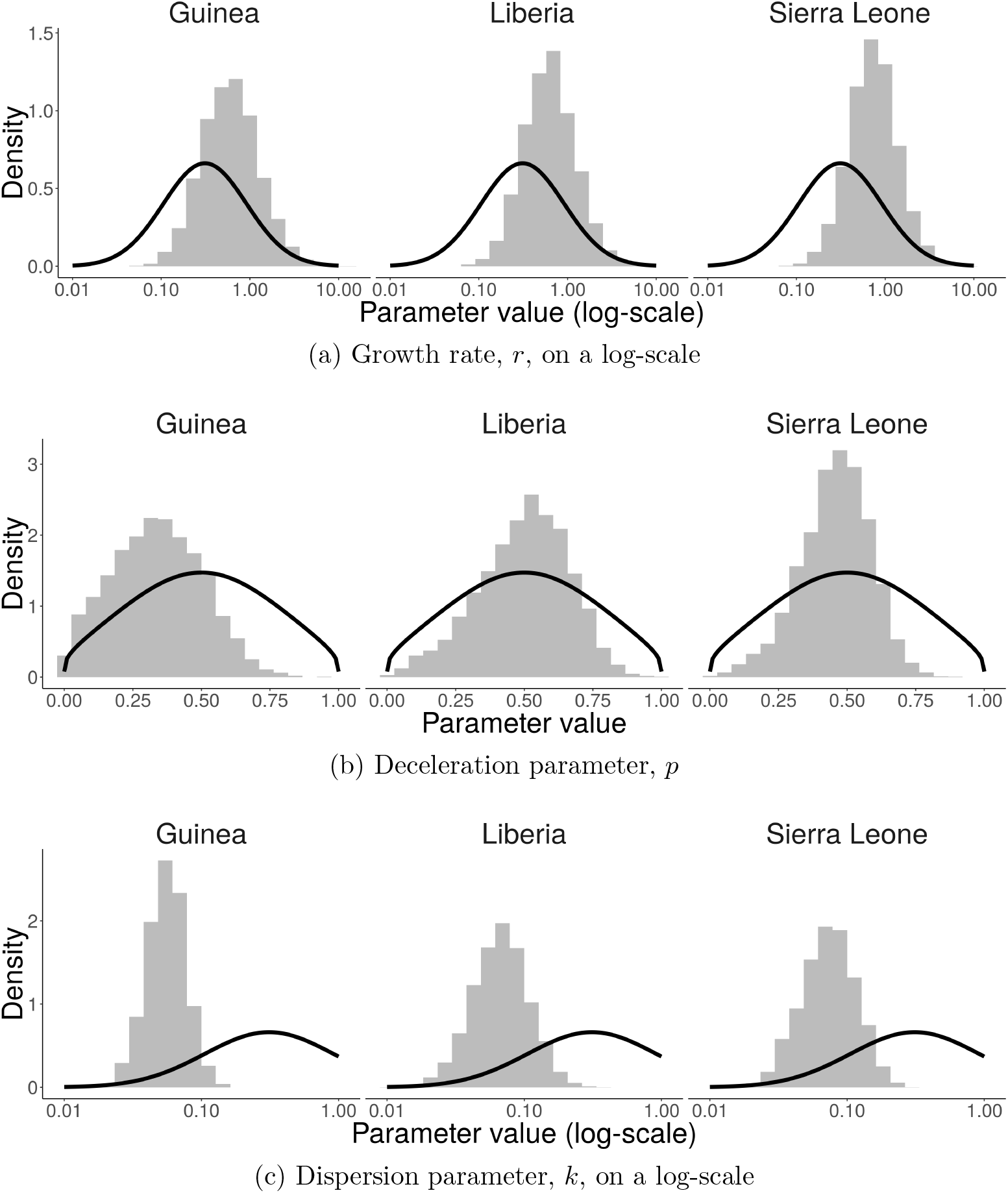
Histograms of posterior samples under the hierarchical model for a.) Guinea, b.) Liberia and c.) Sierra Leone. The marginal prior distribution is included as a solid line to assess convergence.

### 3.2. Comparison of time series and infection chain data from Conakry (Guinea)

Figure 4 shows the marginal posterior distributions for the growth rate, deceleration and dispersion respectively, conditioning upon the time series and secondary infections data from Conakry. The posterior distributions differ from their prior, indicating information was extracted from the data. The parameter estimates inferred from each data set are broadly consistent, suggesting, in this instance, that both data types provide a consistent representation of the dynamics. The time series data suggested a smaller growth rate (mean= 0.38, CI= 0.06 – 0.47) than the tree data (mean= 0.75, CI= 0.17 – 2.52). This trend is reversed for the deceleration parameter, which are 0.30, CI= 0.04 – 0.67 for the time series and 0.13, CI= 0.01 – 0.36 for the tree data. Overall, the time series data suggests slower, but more rapidly accelerating growth than the secondary infections data. We consider potential causes for these differences in the Discussion.

**Fig. 4.**
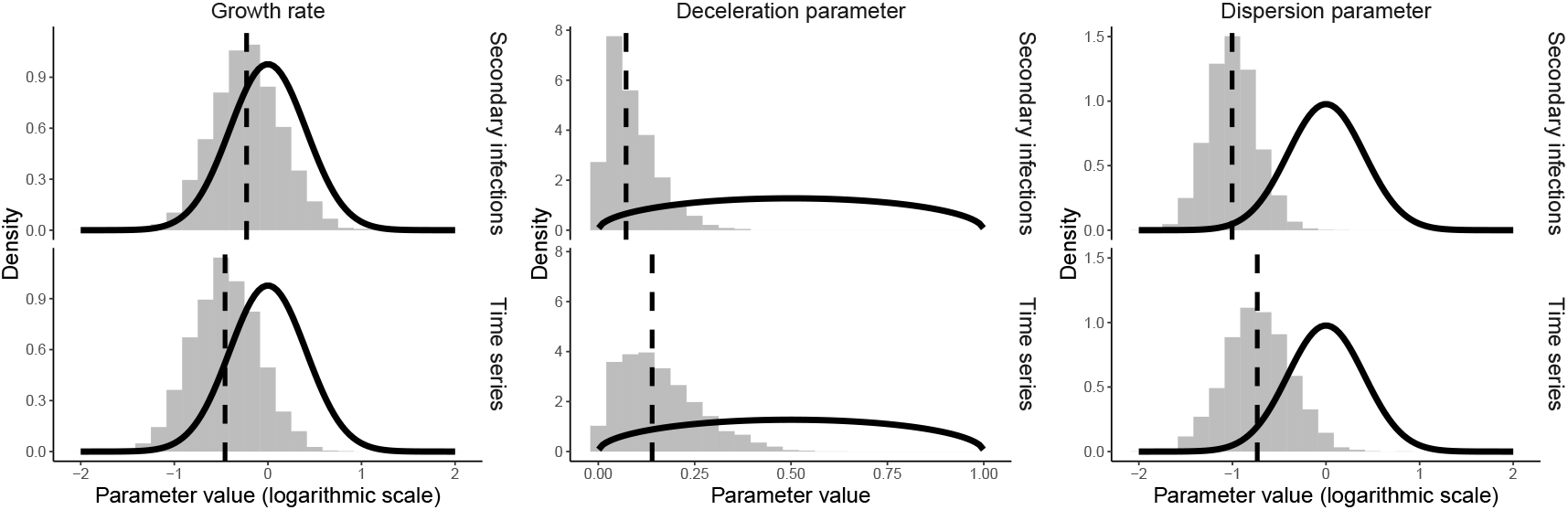
Histograms representing the posterior distribution of the model parameters conditional upon the secondary infections data and the time series data from Conakry. The solid lines show the prior distribution for each of the parameters (obtained via numerical integration). The growth rate and dispersion parameters are shown on a log-scale.

### 3.3. Simulation re-estimation study

Figure 5 shows the simulations of the number of infectious individuals in each generation of the RF-model (Section 2.8). For most of these simulations, the incidence is still increasing during the first 7 generations suggesting the epidemic peak has not yet been reached for the majority of these simulated epidemics.

**Fig. 5.**
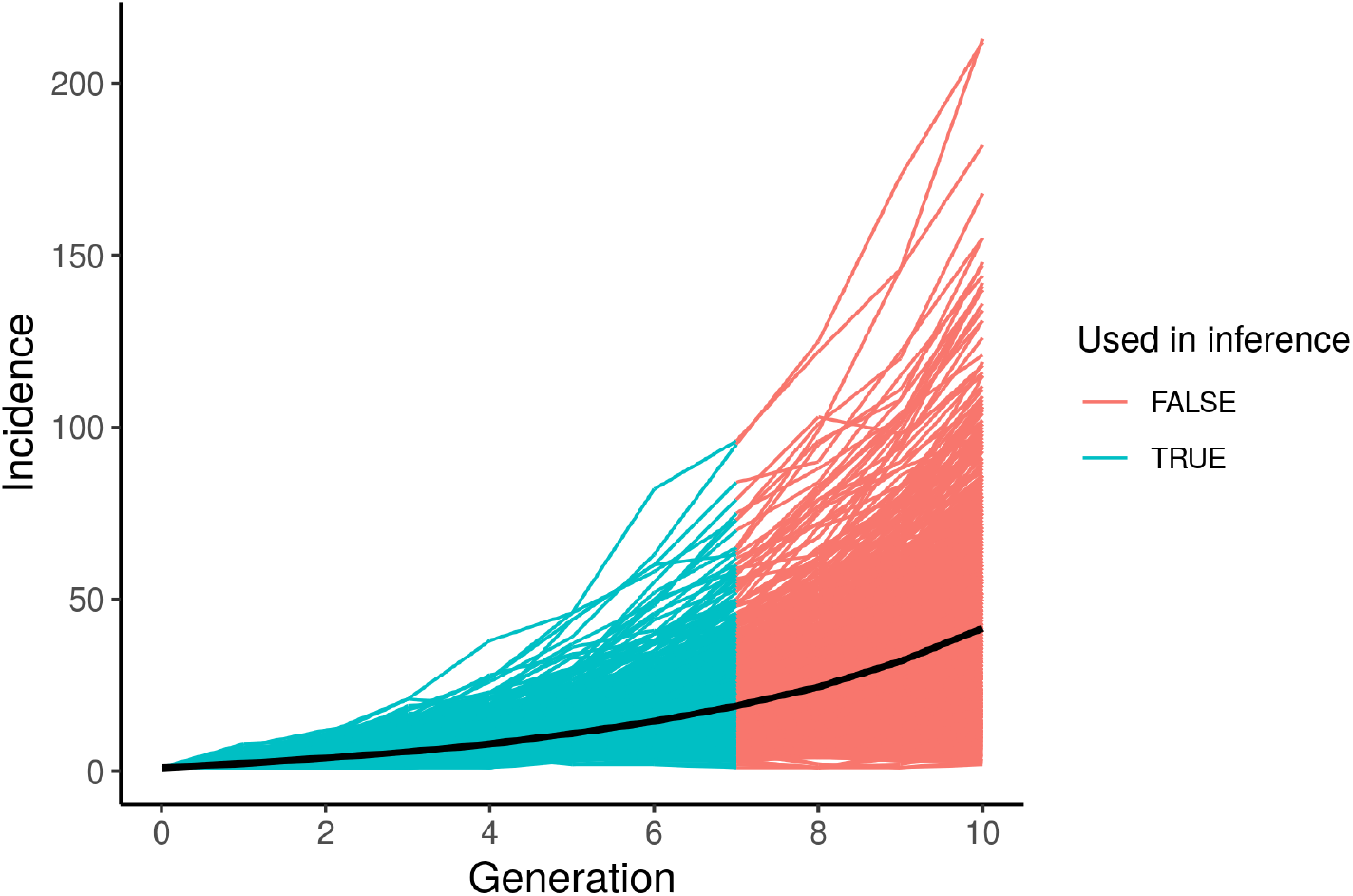
Simulated time series from the Reed-Frost epidemic model and the mean of these time series. The blue portion of the time series was used in the simulation re-estimation study.

Figures 6, 7 and 8 shows the relationships between the maximum a posterior probability (MAP) estimates of the growth rate and the deceleration parameter (respectively) obtained using either data type. In the case of secondary infections data the number of infectious people to “contact trace” is a tuning parameter: a property of the actual observation process. For this study we inspected the number of secondary infections for three different intensities of observations, i.e., we recorded the number of secondary infections from 2, 5 and 10 individuals in each generation.

**Fig. 6.**
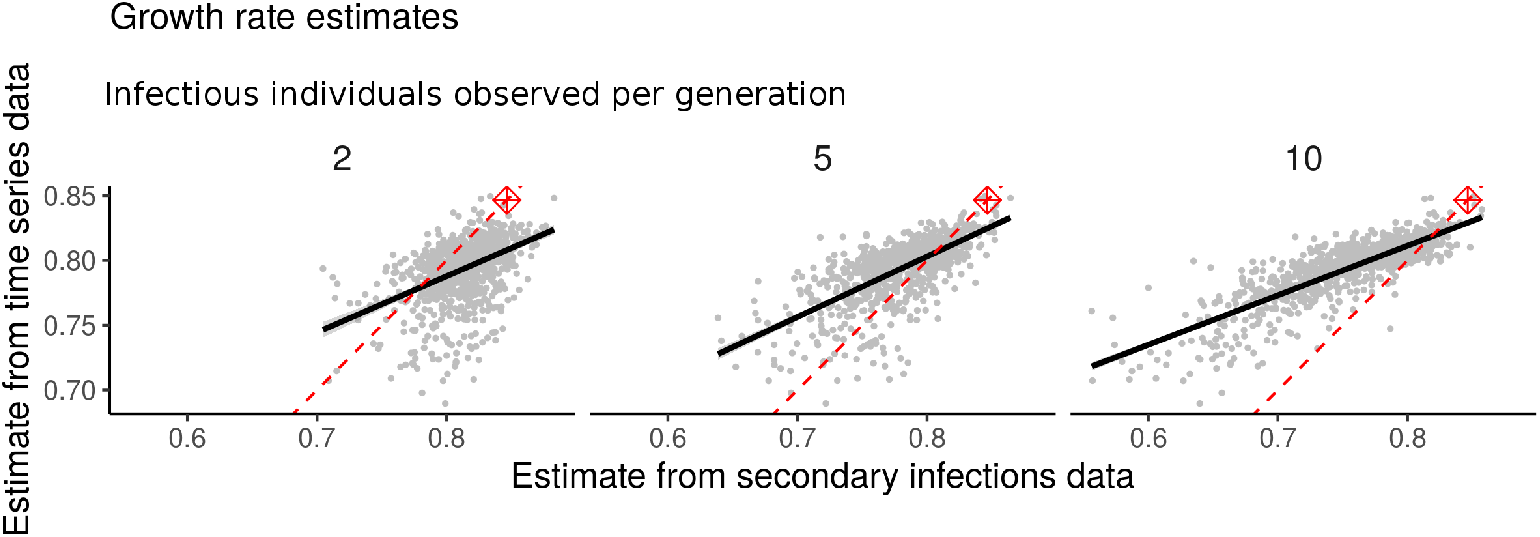
A scatter plot of the maximum posterior probability estimate of the growth rate obtained from the time series data and the secondary infections data. There is a single point for each simulation, and the solid line shows a linear fit with a 95%confidence interval, the dashed line shows the parity line. Each facet shows the estimate conditional upon a different number of observations in each generation: 2, 5, or 10.

**Fig. 7.**
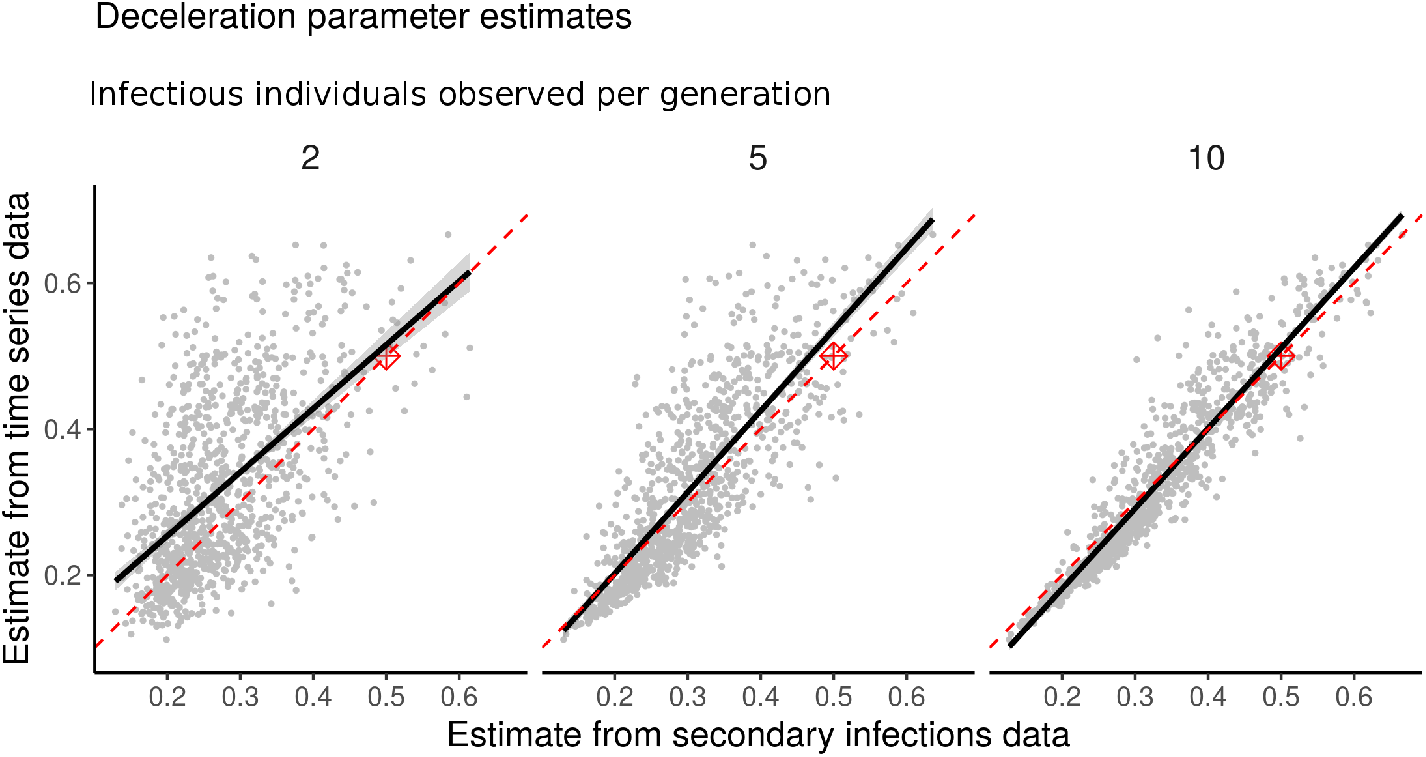
A scatter plot of the maximum posterior probability estimate of the deceleration parameter obtained from the time series data and the secondary infections data. There is a single point for each simulation, and the solid line shows a linear fit with a 95%confidence interval, the dashed line shows the parity line. Each facet shows the estimate conditional upon a different number of observations in each generation: 2, 5, or 10.

**Fig. 8.**
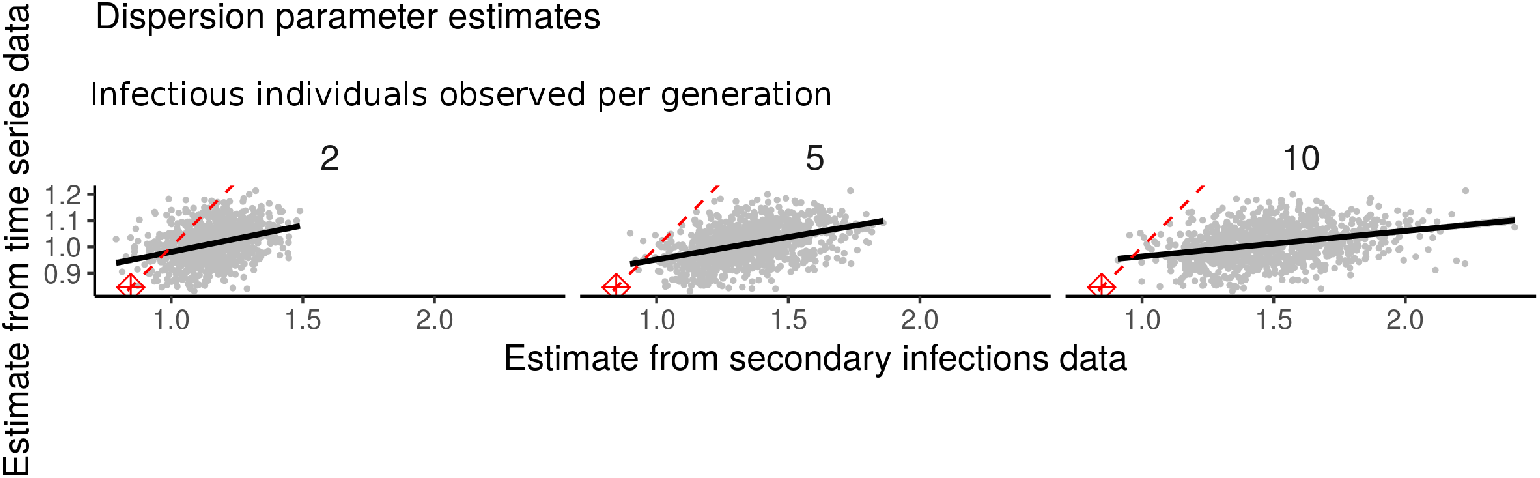
A scatter plot of the maximum posterior probability estimate of the dispersion parameter obtained from the time series data and the secondary infections data. There is a single point for each simulation, and the solid line shows a linear fit with a 95%confidence interval, the dashed line shows the parity line. Each facet shows the estimate conditional upon a different number of observations in each generation: 2, 5, or 10.

Considering the MAP conditional upon each data type, there is a strong correlation between the estimates obtained with each data type for both the growth rate and the deceleration parameter, and this correlation grows stronger as more secondary infections are observed. In the case of the deceleration parameter, once ten individuals have had their secondary infections observed both data types lead to essentially the same estimates. There is a clear bias and increased variability in the estimates derived from the secondary infections data for both the growth rate and the dispersion parameter. As with any Bayesian analysis, it is important to understand the impact of the prior distribution; in the absence of any data, the MAP would be the mode of the prior distribution. In each case, there is a consistent shift in the MAP estimate away from the mode of the prior (as shown in the figures.)

## 4. Interpretation

### 4.1. Sub-exponential growth of EVD in West Africa 2014–2016

The analysis of the EVD time series from Guinea, Liberia and Sierra Leone demonstrates that the in-homogeneous branching process is capable of faithfully describing disease transmission at the population level. The posterior distribution of the deceleration parameter, which controls the scale of the growth, suggests that initially, the incidence grew approximately linearly (and the cumulative incidence quadratically). This differs from the results presented by Chowell *et al* Chowell et al. (2015), who observed that transmission at a sub-national level grew sub-exponential, but that at the national level it grew approximately exponentially. While it is tempting to attribute these differences to the differences in the modelling approach, the most likely explanation is the different pre-processing of the time series data. The previous analysis considered a portion of the time series from later in the epidemic, to mitigate the influence of stochastic effects. Since we have used a stochastic model in this analysis which can explain the initial fluctuations in incidence, we felt it was justified to use data from the start of the epidemic.

### 4.2. Conakry: time series and chains of infection

Using either time series data or secondary infections data from Conakry, Guinea leads to similar parameter estimates demonstrating that either data set could be used to characterise transmission. The time series estimates have a smaller growth rate and a larger deceleration parameter than those from the secondary infections data. The difference in the estimates could be partially attributed to the estimates trading off faster growth (i.e., higher growth rate) for less acceleration of growth (i.e., smaller deceleration parameter.) Since this trade-off should yield similar dynamics over short time spans it is unclear whether this difference would pose substantial issues to interpretation of the parameters.

In the case of this Ebola epidemic, the time series data was available long before the infection tree. However, obtaining comprehensive time series of disease is challenging, and it is interesting to know that there are alternative data sets which may be useful and already part of the data collected during intervention measures such as contact tracing. Moreover, our observations do not guarantee that we can rely on the agreement between the inference methods in general which is, in part, why we also carried out the simulation study.

### 4.3. Simulation re-estimation study

The simulations from the Reed-Frost (RF) model (shown in Figure 5) emphasise the variability between realisations of stochastic epidemic models, and consequently, the substantial role stochasticity plays during outbreaks. The parameter estimates derived from the time series data and secondary infections data generated by these epidemics have a strong correlation which increases with the number of secondary infections observed. However, for the growth rate there is a clear trend that the secondary infections data tends to yield lower point estimates for the growth rate. A difference of this kind should not be ignored, however, given there will also be a level of uncertainty on these estimates, they will still give broadly consistent characterisations of the epidemic. The simulations used where generated with an RF model so there is not an obvious ground truth to compare these values to in order to further investigate which of the estimators is biased.

Together, this demonstrates that characterisations derived from each data type will be similar given a sufficient number of secondary observations, however (particularly in the case of the growth rate) there are systematic differences in the estimates that we were unable to explain. Conditioning the process against extinction in the case of the time series estimator, but not in the case of the secondary infections estimator, may be contributing to this systematic difference.

## 5. Discussion

We have presented an in-homogeneous branching process to model outbreaks of a transmissible pathogen. The simplicity of the process means we can construct both a population and an individual scale view and subsequently assimilate data from either scale.

Our model admits a closed form for the likelihood for the time series and we have supplied an approximation for the likelihood of the secondary infections data. These closed forms make it feasible to conduct a Bayesian analysis and handle subtleties of the fitting process (in the case of the time series data), such as conditioning the process against extinction to account for the implicit observation bias. While we do not address unobserved cases in the secondary infections data, nor do we have a sophisticated method for aggregating cases into generations, our analysis of the Ebola data suggests these limitations do not cause substantial problems with the inference.

From that Ebola data, our analysis provides clear evidence for sub-exponential growth and a significant role for stochasticity in shaping the early epidemic dynamics. Our analysis extends work carried out during the 2014 Ebola epidemic by Chowell *et al* Chowell et al. (2015) and a comprehensive study of the dynamics of several pathogens’ transmission Viboud et al. (2016). We used the same phenomenological model as the phenomenological backbone of the branching process. The resulting process has the same dynamics (on average) but with a mechanistic underpinning. This enables us to handle a wider range of data types, for example, the tree data from Faye *et al* Faye et al.(2015).

Assimilating secondary infection and time series data types simultaneously was beyond the scope of the current work. Since the conditional distribution of Poisson variables given their sum is multinomial, it would be feasible to perform simultaneous assimilation with a Poisson offspring distribution. However, the matter becomes more complicated when using a negative binomial distribution, as we use here due to the over-dispersion so common in epidemiological transmission data.

From the simulation study, we have identified a systematic difference in the estimators for model paramters based on the two data types. We suspect this stems from thediffering treatment of extinction for the two analyses. A more in-depth study of this was beyond the scope of the current work.

Lags in time series data cause substantial problems when forecasting incidence Moss et al. (2018). “First Few Hundred” (FF100) studies collect the same type of data as contact tracing and are heralded as a way to rapidly provide a characterisation of transmission dynamics Black et al. (2017). While for the 2014 Ebola epidemic the time series was available before the secondary infections tree, there does not seem to be anything intrinsic to the data collection process that precludes this being reversed. In fact, it seems plausible that in active surveillance programs and with increased use of sequencing, secondary infections data may become available before time series, and so the method we present here may have important application in future real-time outbreak analyses. Of course, there are ethical, procedural, and technical challenges that are introduced by collecting, analysing and storing data such as this since by its very nature it resolves more of the epidemic. The source code to carry out the analyses reported in this paper are publically available under an open-source licence at https://bitbucket.org/azarebski/subexp.

Most pertinent to improving the value of our approach is establishing how to handle incomplete secondary infections data. We investigated the consequences of partial observation of the infectious population, but with perfect ascertainment of the number of infections due to each individual. A natural extension then is to consider partial observation of the population with imperfect resolution, i.e., observe a random subset of the infectious population and observe only a subset of their infections. This additional way in which data can be missing is particularly important in airborne disease, such as influenza, where the source of an infection may be harder to ascertain. If the goal is to characterise the transmission dynamics of a pathogen for which sub-clinical cases are rare, such as Ebola virus disease, then the assumption of complete observation among those observed does not seem unreasonable. As sequencing data becomes more readily available we will have improved capability to determine who-infected-whom and models such as the one presented in this work are poised to take advantage of this additional information.

## 6. Acknowledgements

The authors acknowledge the helpful discussions with Peter Dawson, Gerardo Chowell and Peter Taylor during the conceptualisation of this work. AEZ was supported by an Australian Government Research Training Program scholarship while undertaking his PhD, with further support from an Australian Government Defence Science Technology Group scholarship. RM was supported by an Australian Government Defence Science Technology Group research agreement.

